# Substrate, temperature, and geographical patterns among nearly 2,000 natural yeast isolates

**DOI:** 10.1101/2021.07.13.452236

**Authors:** William J. Spurley, Kaitlin J. Fisher, Quinn K. Langdon, Kelly V. Buh, Martin Jarzyna, Max A. B. Haase, Kayla Sylvester, Ryan V. Moriarty, Daniel Rodriguez, Angela Sheddan, Sarah Wright, Lisa Sorlie, Amanda Beth Hulfachor, Dana A. Opulente, Chris Todd Hittinger

## Abstract

Yeasts have broad importance as industrially and clinically relevant microbes and as powerful models for fundamental research, but we are only beginning to understand the roles yeasts play in natural ecosystems. Yeast ecology is often more difficult to study compared to other, more abundant microbes, but growing collections of natural yeast isolates are beginning to shed light on fundamental ecological questions. Here we used environmental sampling and isolation to assemble a dataset of 1,962 isolates collected from throughout the contiguous United States of America (USA) and Alaska, which were then used to uncover geographic patterns, along with substrate and temperature associations among yeast taxa. We found some taxa, including the common yeasts *Torulaspora delbrueckii* and *Saccharomyces paradoxus*, to be repeatedly isolated from multiple sampled regions of the US, and we classify these as broadly distributed cosmopolitan yeasts. A number of yeast taxon - substrate associations were identified, some of which were novel and some of which support previously reported associations. Further, we found a strong effect of isolation temperature on the phyla of yeasts recovered, as well as for many species. We speculate that substrate and isolation temperature associations reflect the ecological diversity of and niche partitioning by yeast taxa.

**Take Away:**

- Analysis of environmental metadata of nearly 2,000 yeast isolates.
- Individual yeast taxa associate with specific substrates and plant genera.
- Optimal yeast isolation temperature differs depending on taxonomic rank.
- Substrate type and isolation temperatures affect isolated yeast diversity.

## Introduction

Yeasts are fungi that spend at least part of their life cycle in a unicellular state and do not form fruiting bodies (Kurtzman, Fell and Boekout 2011). They are globally distributed, speciose, and diverse, and many have important relationships with humans. Yeasts are polyphyletic and are represented in the phyla Ascomycota (subphyla Saccharomycotina and Taphrinomycotina) and Basidiomycota (Nagy et al. 2014). The ascomycete subphylum Pezizomycotina, which is sister to the Saccharomycotina, also includes some dimorphic fungi that spend significant portions of their life cycle in a unicellular state, but they are not traditionally regarded as yeasts (Kurtzman, Fell and Boekout 2011). The phylogenetic, phenotypic, and genomic diversity within yeasts suggests that divergent yeasts have distinct ecological roles. Despite their importance and ubiquity, the ecology and distribution of most yeast species remain poorly understood. The advances in culture-independent metagenomic characterization of microbial communities that have enabled increasingly detailed characterization of bacterial communities (Franzosa et al. 2015) do not lend themselves well to the study of yeasts; typical methods for environmental detection of bacterial microbes are less useful for eukaryotic microbes that are low-abundance community members (Pereira-Marques et al. 2019). As a result, most research in yeast ecology is reliant on environmental sampling and isolation in synthetic media, either by direct plating (Boynton, Kowallik, Landermann and Stukenbrock 2019; Rose, Winston and Hieter 1990; Stefanini et al. 2012) or enrichment (Knight and Goddard 2015; Liti, Warringer and Blomberg 2017; Sniegowski, Dombrowski and Fingerman 2002).

Isolation-dependent approaches are admittedly not ideal for studying ecology because most investigations are performed away from the sample collection site in laboratory conditions that differ drastically from natural conditions, because of the presence of bottlenecks in the isolation process, and because most fungal diversity is likely unculturable (James, Stajich, Hittinger and Rokas 2020). Despite these limitations, a broad and dense enough sampling strategy can begin to illuminate the cryptic ecology of yeasts. For example, exhaustive environmental sampling revealed that *S. cerevisiae* is not the purely domestic species it was once believed to be (Török, Mortimer, Romano, Suzzi and Polsinelli 1996; Vaughan-Martini and Martini 1995), which has led to a much better understanding of the global distribution of *S. cerevisiae* (Naumov, Naumova and Sniegowski 1998; Peter et al. 2018; Wang, Liu, Liti, Wang and Bai 2012). Similar insights have been gained into the distributions of other members of the genus *Saccharomyces* through isolation-based surveys (Charron, Leducq, Bertin, Dubé and Landry 2014; Langdon et al. 2020; Nespolo et al 2020). While studies, such as these, are beginning to elucidate the distributions (and in some cases ancestral origins of) well-studied species, there is still much to be understood about the natural habitats and ecology of the vast majority of species.

Most surveys of the natural habitat of yeasts are limited in sampling depth and the breadth of substrate types sampled and therefore lack statistical power to assign preferred substrates to yeast taxa. Recent analyses that have overcome these limitations by using large datasets of isolates have been able to uncover associations between yeast species and habitats (Charron et al. 2014; Opulente et al. 2019; Sylvester et al. 2015). Connecting yeast species to biotic or abiotic substrates is the first step in inferring specific niches. Hypotheses regarding the ecological roles of yeasts in nutrient cycling, symbioses, and microbial interactions can be generated and tested once their preferred substrates are known. These connections also provide practical information about the natural reservoirs of yeasts that are significant to humans, both those that pose threats as emerging pathogens (Bensasson et al. 2019; Friedman and Schwartz 2019; Geddes-McAlister and Shapiro 2019; Opulente et al. 2019) and those that are potentially valuable for the development of bioindustrial capabilities (Cordente, Schmidt, Beltran, Torija and Curtin 2019; Hittinger, Steele and Ryder 2018; Spagnuolo, Wohlbach et al. 2011, Yaguchi and Blenner 2019).

The most direct use of the characterizations of the natural reservoirs of different yeast taxa would be of applied use in the pragmatic design of further isolation protocols. Individual species or groups of species could be targeted by deep sampling of associated substrates. Likewise, surveys of yeast communities would benefit from sampling substrates that are known to yield high levels of diversity. Yeasts have historically been isolated mostly from fruit, soil, and bark, but there have been successful isolations from many other substrates, including aquatic environments, flowers, leaves, and insects (Han et a., 2015; Roth Jr, Orpurt and Ahearn 1964; Sláviková, Vadkertiová and Vránová 2007; Stefanini et al. 2016). Coupling diversity of substrates with regional distribution of yeasts could lead to more precision in isolation strategies.

In an effort to better understand the distribution and natural ecology of yeasts, we have launched an ongoing initiative to sample yeast diversity from natural substrates. Previously, our lab investigated preferences and associations of yeasts among 589 isolates (Sylvester et a., 2015). This growing collection has also described associations of pathogenic yeasts and their geographic distribution in the United States of America (USA) (Opulente et al. 2019). Here, we have expanded this collection to curate a dataset of 1,926 isolates. To our knowledge, the collection analyzed here is the largest collection of natural yeast isolates gathered using common isolation protocols that has been published to date. We used this dataset to examine the spatial distribution patterns of diverse yeast species, look for associations between specific taxa and habitats, and examine overall yeast diversity among different types of sampled substrates. Using spatial and temporal isolation patterns, we also identified yeasts that exhibited a cosmopolitan distribution within our dataset. Due to the statistical power afforded by such a large dataset, we confirmed previously suggested substrate associations and identified new ones. Finally, we examined the diversity of yeasts associated with different substrate types to better understand where yeast communities are expected to be especially complex.

## Methods

### Sample collection and yeast isolation

Samples of natural substrates were collected across the continental USA and Alaska. Data recording the GPS location; substrate types; and, if applicable, plant species accompanied all collections. Sampling sites were generally natural areas, and sites that were strongly associated with humans or industry were avoided. Samples were collected using a sterile bag and without any human contact to prevent contamination. These samples were then either processed immediately or stored at 4°C for up to three weeks. Isolation of yeasts from samples was performed as described in Sylvester et a., 2015, with the addition of 4°C as an isolation temperature. Briefly, samples were inoculated in a 15-ml conical tube into 9ml of Wild Yeast Medium (5 g/L (NH_4_)_2_S0_4_, 1 g/L Synthetic Complete Dropout mix (US Biological), 1.72 g/L YNB w/o AA, Carbohydrate & w/o AS (US Biological), 0.1 g/L Ampicillin, 0.03 g/L Chloramphenicol), supplemented with either 8% or 0.8% glucose. The Sylvester et al. 2015 recipe included (NH_4_)_2_S0_4_, but it was mistakenly excluded from the Materials and Methods. Inoculated tubes were then incubated at one of four temperatures, 4°C, 10°C, 22°C, or 30°C, until there was evidence of growth or fermentation (generally 1-2 days at warm temperatures, and up to two months at cold temperatures). Cultures were then passaged once more through 4 ml Wild Yeast Medium in a 5-ml conical tube and incubated at their respective temperatures until growth was evident. Enriched cultures were plated to YPD agar plates. Plates were inspected for discrete colonies with distinct morphotypes. Colonies with spreading hyphal morphologies were typically avoided. This selection filtered isolates based on colony morphology, such that some Pezizomycotina species that form characteristic “yeast-like” colonies were occasionally isolated, while most Pezizomycotina species were not. Single colonies were selected for identification, restruck to purify, and frozen down in 15% glycerol stocks.

Taxonomy was assigned through amplification and sequencing of the ITS region of the *rDNA* locus. Specifically, the entire ITS region was amplified using the oligonucleotides ITS1 and NL4 (McCullough, Clemons, McCusker and Stevens 1998; O’Donnell 1993), and the ITS1/2 region was sequenced using the the oligonucleotide ITS4 (McCullough et al. 1998). ITS sequences were queried against the nucleotide database of GenBank using BLASTN, and query sequences that matched their best database hit by 97% or more were positively identified based on that search. Queries that did not return matches of at least 97% were sequenced at the D1/D2 domain of the *rDNA* locus using NL4 as a sequencing oligonucleotide to achieve a positive identification. Isolates that were more than 3% divergent in the D1/D2 domain were considered candidates for novel species, which will be formally described elsewhere. Forty-six such isolates were identified to the level of 18 different genera, and all potentially novel species are indicated in **Suppl. Table 1**. Some isolates can only be identified to the level of species complex due to poorly delineated species phylogenies. Because there are a number of candidate novel species and isolates belonging to species complexes, we use the term operational taxonomic unit (OTU) in lieu of species throughout.

A detailed protocol is provided in **Suppl. Note 1**.

### Dataset curation

In some instances, we were unable to amplify the ITS or D1/D2 sequences of an isolate. Any strain that fell into this category was subject to three independent attempts to achieve a sequence. If the isolate failed three times, they were excluded from analyses.

Oftentimes, multiple isolates of the same OTU isolated from the same exact substrate sample appear in the dataset. To eliminate these possibly duplicate strains, prior to all analyses, except one category of analysis as described below, identical OTUs isolated from the same processed sample were eliminated. Exclusion of identical OTUs isolated from the same processed sample resulted in 1,518 unique isolations (**Fig. S1**). The same environmental sample was sometimes subjected to multiple different temperatures for isolation, and these were considered separate isolations for temperature analyses only.

### Detection of cosmopolitan OTUs

We sought to use our extensive dataset of isolation locations to determine which OTUs in our data could be considered cosmopolitan across the expansive sampled region. Instead of simply determining which OTUs were frequently isolated temporally or spatially, we considered differences in sampling depth across the USA to identify those OTUs that were always isolated in regions that were deemed sufficiently sampled to detect them. We began by dividing the country into ten regions, the Arctic (Alaska) plus the nine climatic regions of the contiguous USA defined by the National Oceanic and Atmospheric Association (NOAA) (**Fig. S2**). For each OTU, we then calculated an expected discovery rate based on the most densely sampled climatic region in which that OTU was found. For example, *Torulaspora delbrueckii* was isolated 60 independent times out of 382 total samples collected in the Upper Midwest, giving it an isolation rate of once every 6.3 samples (382 total samples/60 OTU isolations). Thus, we would then expect *T. delbrueckii* to be isolated in all regions that had been sampled at least 7 times. In this manner, we determined the regions from which each OTU was expected to have been isolated and compared these to the regions in which each OTU was actually found. We eliminated from consideration all yeasts that had only been isolated only once (116 OTUs) and all those yeasts that were only expected in a single region (80 OTUs, all expected only in the Upper Midwest due to high sampling density). We defined cosmopolitan yeasts as OTUs that were isolated either in all regions where they were expected or in all but one region where they were expected. For example, *T. delbrueckii* was expected in all regions except the Northern Rockies and Southwest, and it was found in all expected regions with the single exception of the Arctic. Since we allowed for one region of not being detected where expected, *T. delbrueckii* was therefore categorized as a cosmopolitan yeast OTU.

### Association analyses

Enrichments of yeast taxonomic groups (phyla and subphyla classifications) in substrate-type categories and isolation temperatures were examined with one-tailed Fisher’s exact tests followed by Benjamini Hochberg post-hoc corrections. Positive associations at the level of individual yeast OTUs were detected by independently permuting OTU-substrate, OTU-substrate genus, or OTU-temperature combinations to find those combinations present more often than expected by chance. We did not examine our data for depleted combinations because we cannot statistically distinguish between true depletion and undersampling. P-values and expected rates were calculated based on 10,000 permutations for each association analysis. Benjamini-Hochberg adjustments were applied to P-values. Permutations were performed by completely resampling yeast OTUs assigned to each substrate or temperature category without replacement using custom R scripts.

### Diversity analyses

All diversity analyses were carried out using the R vegan package (Oksanen et al. 2019). We used both rarefaction and Shannon entropy indices (H’) to compare alpha diversity among substrate types and isolation temperatures. Because comparison of H’ indices are complicated by uneven sampling, we used rarefaction curves to corroborate differences in sampled diversity. For substrate alpha diversity, we analyzed only specific substrate types sampled more than five independent times. All four isolation temperatures were included in temperature alpha diversity analyses. Beta diversity among isolation temperatures was estimated by calculating the Jaccard distance (*D*_*J*_).

## Results & Discussion

### Curated dataset of isolates

The final dataset curated for this study consisted of 1,962 isolations of 262 unique operational taxonomic units (OTUs) isolated from 688 unique samples (**Suppl. Table 1**). Rarefaction indicated that the true richness is not represented in the sampled range and that this sample is an underestimate of richness (**Fig. S3A**). 483 strains in this study were also included in a previously published survey of wild yeast isolates (Sylvester et al. 2015), as were an additional 34 strains from subsequent taxon-specific studies (Krause et al. 2018; Langdon et al. 2020; Leducq et al. 2014; Opulente et al. 2019; Peris et al. 2016; **Suppl. Table 1**). 106 strains from Sylvester et al. 2015 were excluded due to differences between isolation protocols, missing metadata, or other discrepancies that prevented inclusion in the present analyses. Accompanying metadata for each isolate included GPS coordinates of sample collection, the general substrate type, the specific substrate type, the genus and species of biotic substrates, the incubation temperature used in laboratory isolation, and the ITS sequences used to identify the species. Yeasts from both major divisions or phyla of Dikarya, Basidiomycota (90 OTUs) and Ascomycota (172 OTUs), were represented in the data. Within Ascomycota, isolates belonged to the subphyla Saccharomycotina (153 OTUs), Pezizomycotina (13 OTUs), and Taphrinomycotina (1 OTU) (**Fig. S3B, Fig. S4**). Pezizomycotina and Taphrinomycotina were represented by relatively few isolates due to systematic avoidance of colonies belonging to the former (see methods) and the presumed low abundance and need for specialized enrichment for the latter (Benito, Calderón and Benito et al. 2018).

### Geospatial variation among isolations

#### The Upper Midwest was overrepresented in our dataset

Isolates were obtained from all major climate regions of the contiguous United States and Alaska, except for the Southwest (**Fig. S2**). The Upper Midwest was the most densely sampled region with 868 unique isolations from 382 samples, followed by 144 isolations from 53 samples in the Ohio Valley, 140 isolations from 58 samples in the Northeast, 136 isolations from 87 samples in the Northwest, 105 isolations from 55 samples in the Southeast, 40 isolations from 14 samples in the West, 32 isolations from 16 samples in the South, and 6 isolations from 3 samples in the Northern Rockies. Zero isolates were collected from the Southwest. An additional 44 isolations came from 31 Alaskan samples, which we classified as Arctic.

#### Rarely and frequently isolated OTUs

Of the 262 unique OTUs, 116 were singletons that were found in only one isolation (**Suppl. Table 2**), while the remaining 146 generally were isolated fewer than thirty times. The singleton OTUs isolated in this study were primarily isolated from bark and soil, but singletons were not enriched for any substrate category (**Fig. S5A**). Singletons were, however, enriched for yeasts belonging to the Basidiomycota (*P*_adj_= 0.003) and Pezizomycotina (*P*_adj_= 0.0005) (**Fig. S5B**), reflecting an isolation protocol designed to capture Saccharomycotina species. Singletons were evenly distributed across sampled regions (**Fig. S5C**).

Sixteen yeast OTUs were isolated over 20 independent times in this dataset (**Suppl. Table 3)**. Among these were two *Saccharomyces* species, *Saccharomyces paradoxus* and *Saccharomyces cerevisiae*, for which considerable population genetic work has already been done (Xia et al. 2017; Peter et al. 2018, Hénault et al. 2017). Other frequently isolated OTUs are of broad interest but have been subjected to relatively little population genetic analysis. These include *Kluyveromyces lactis, T. delbrueckii, Hanseniaspora uvarum*, and the *Metschnikowia pulcherrima* spp. complex. The collection described herein is teeming with potential for population-level studies for these and more OTUs.

#### 11 cosmopolitan OTUs

Although many yeast species are frequently referred to as cosmopolitan, there are no clear criteria by which yeasts are or are not classified as such. We examined our dataset for yeasts that were widely distributed across the USA by considering both the number and spatial distribution of isolations for each OTUs. Briefly, we defined an OTU-specific isolation rate and then looked for OTUs that were generally able to be isolated in all regions where sampling was dense enough to detect them (**Suppl. Table 4**, see Methods). While this approach cannot exclude any yeasts from being broadly distributed, it can identify those OTUs in our dataset that are most likely cosmopolitan in distribution. We determined that 11 OTUs in our dataset were cosmopolitan and generally found when expected across climatic regions (*Candida railensis, Candida sake, Cryptococcus flavescens, Cyberlindnera saturnus, Debaryomyces hansenii, Leucosporidium scottii, Rhodotorula fujisanensis, S. paradoxus, Scheffersomyces ergatensis, T. delbrueckii*, and *Wickerhamomyces anomalus*, **Suppl. Table 5**). Only 5 of these OTUs were also among the 16 OTUs isolated over 20 times, which suggests that frequent isolation in individual regions may not indicate otherwise broad geographic distributions. Three of the OTUs we identified as cosmopolitan were isolated less than 10 times (*Debaryomyces hansenii, Leucosporidium scottii, Rhodotorula fujisanensis*), but were nonetheless always detected in regions with sufficient sampling (**Fig. S6A, Fig. S7C**). Cosmopolitan OTUs ranged from being isolated in just two regions (due to requiring sufficiently dense sampling to detect them) to being isolated in a maximum of 7 regions (**Fig. S6B, Fig. S7C**). No OTU was isolated from all nine sampled USA regions. Cosmopolitan yeasts were not isolated at equal rates among substrates, but rather were significantly enriched for soil samples (*P*_adj_= 3.09E-05, **Fig. S7A**).

Our approach to defining cosmopolitan isolates is imperfect and affected by sampling biases in our data, but the clear statistical criteria represent an improvement over the ad hoc subjective assessments that are sometimes made. The approach only considered total climatic region sampling density and not the sampling densities of any specific substrates. Most substrates were not uniformly sampled across regions, and yeasts with high substrate-specificities may have been missed. Unfortunately, powerful approaches for determining distributions for yeast OTUs remain elusive. They are often not abundant enough to be detected by metagenome sequencing or to be reliably isolated in all samples. The clandestine biogeography of yeasts is mainly studied using the same approaches we use here – laborious environmental sampling and laboratory isolation. While it is possible that many more yeast OTUs in our collection may also have wide distributions, we can at least conclude that the eleven OTUs we identify as cosmopolitan show patterns of ubiquitous distributions, at least across the USA. Soil was one of the most broadly sampled substrate types across climatic regions, which may be one reason the OTUs we identify as cosmopolitan seem to be soil-associated taxa. Conversely, soil is a ubiquitous substrate, and the strong statistical enrichment for cosmopolitan yeasts among soil samples may suggest soil as a reservoir of cosmopolitan yeasts.

### Substrate and temperature associations

#### Substrate – higher taxonomic group associations

We were able to assign one of 40 specific substrate categories to 1,522 isolations, which had yielded 161 unique yeast OTUs (**Fig. S8A**). Contingency tables were used to identify associations between higher taxonomic groups of yeast (phylum Basidiomycota and the three Ascomycota subphyla) and substrate. When taxonomic group representation across substrate types was examined (**Suppl. Table 6**), we found that leaves were enriched for Basidiomycota (*P*_adj_= 0.017) but found no other significant enrichment of taxonomic groups across substrate types.

#### Individual OTU - substrate associations

Permutations were used to identify associations between specific taxonomic units and sampled substrates that occurred more often than expected by chance. Of the 714 observed combinations (**Fig. S9A**), we found 23 yeast OTU-by-substrate associations that occur more frequently than expected by chance (*P*_adj_< 0.05, **Fig. 1A, Suppl. Table 7**). Soil, which was the most heavily sampled substrate, was significantly associated with two Basidiomycota and three Saccharomycotina OTUs (*Mrakia* spp. complex, *P*_adj_= 0.01; *Trichosporon porosum, P*_adj_= 0.032; *T. delbrueckii, P*_adj_= 0.014; *Cyberlindnera saturnus, P*_adj_= 0.014; *S. paradoxus, P*_adj_= 0.022). Bark was also heavily sampled but yielded only two significant associated OTUs, both of which were Saccharomycotina (*Lachancea kluyveri, P*_adj_= 0.034; *Sc. ergatensis, P*_adj_= 0.032). Leaf samples were also associated with two OTUs, both of which were Basidiomycota (*Rhodotorula nothofagi, P*_adj_= 0.022; *Mrakia gelida, P*_adj_= 0.022). Fungal samples were significantly associated with three Saccharomycotina OTUs (*H. uvarum, Suhomyces bolitotheri, Teunomyces cretensis/kruisii* spp. complex, *P*_adj_< 0.0001); fruit with one Basidiomycota OTU and two Saccharomycotina OTU (*Curvibasidium cygneicollum, P*_adj_= 0.019; *Pichia kudriavzevii, P*_adj_= 0.022; *H. uvarum, P*_adj_= 0.034; respectively); sand with two Saccharomycotina OTUs (*Kazachstania serrabonitensis, P*_adj_< 0.0001; *Pichia scaptomyzae, P*_adj_= 0.022). Feathers, flowers, insects, lichens, and needles were associated with a single OTU each (*Peterozyma toletana, P*_adj_= 0.034; *Zygowilliopsis californica, P*_adj_< 0.0001; *Kwoniella newhampshirensis, P*_adj_= 0.029; *Scleroconidioma sphagnicola, P*_adj_= 0.01; *Sydowia polyspora, P*_adj_*=* 0.032; respectively). Plant matter samples (excluding matter that fell into other plant-related categories) were associated with a single OTU (*Candida mycetangii, P*_adj_= 0.014).

**Figure 1.**
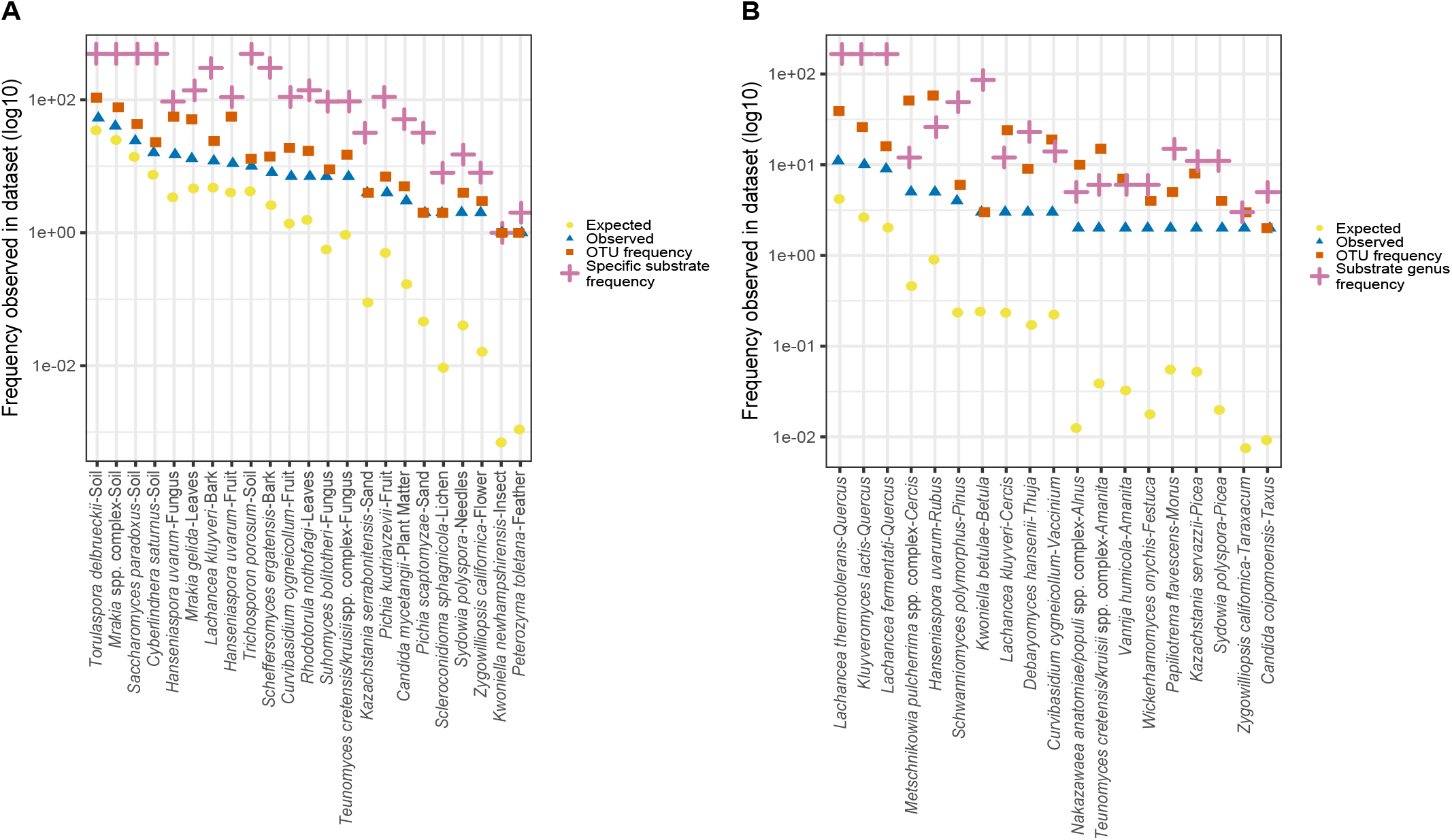
A) Permutations identified 23 yeast OTU – substrate category associations observed more often than expected (*P*_adj_<0.05). Blue triangles indicate the number of times each combination was observed, red squares indicate the number of times the yeast OTU was observed in the permuted dataset, yellow circles indicate the expected rates of each association, and pink plus signs indicate the number of times the substrate category was observed in the permuted dataset. Associations are ordered by the frequency of the observed combinations. **B)** Permutations identified 19 yeast OTU – substrate genus associations observed more often than expected (*P*_adj_<0.05). Blue triangles indicate the number of times each combination was observed, red squares indicate the number of times the yeast OTU was observed in the permuted dataset, yellow circles indicate the expected values of each association, and pink plus signs indicate the number of times the substrate genus was observed in the permuted dataset. Associations are ordered by the frequency of the observed combinations.

When possible, substrate genera were identified for plant and fungal substrates directly sampled (**Fig. S8B)**. Substrate genera were also assigned to samples indirectly associated with plant genera (e.g. soil sampled from base of tree). In total, 66 substrate genera were identified across 1,026 isolations comprising 209 OTUs (**Fig. S9B, Suppl. Table 8**). Nineteen of 658 observed substrate genera – yeast associations were found more often than expected by chance (**Fig. 1B**). *Quercus*, the genus to which oak trees belong, was heavily sampled due to reported associations with *Saccharomyces* species (Naumov 1996; Naumov, Naumova and Sniegowski 1998; Sniegowski et al. 2002; Kowallik and Greig 2016; Wang et al. 2012; Zhang, Skelton, Garner and Goddard 2010). Three Saccharomycotina OTUs were significantly associated with *Quercus* spp., (*Lachancea fermentati, P*_adj_< 0.0001; *Kluyveromyces lactis, P*_adj_ = 0.008; *Lachancea thermotolerans, P*_adj_ = 0.035). Notably, none of the *Quercus*-associated OTUs belong to the genus *Saccharomyces*, although all three do belong to the Saccharomycetaceae, the family to which *Saccharomyces* belongs. Spruce trees of the genus *Picea* were associated with the dimorphic Pezizomycotina OTU *Sy. polyspora* (*P*_adj_*=* 0.008) and the Saccharomycotina OTU *Kazachstania servazzii* (*P*_adj_ = 0.035). *Cercis*, a genus of large flowering shrubs, was associated with two Saccharomycotina OTUs, the *M. pulcherrima* spp. complex (*P*_adj_<0.0001) and *L. kluyveri* (*P*_adj_= 0.048). *Amanita* was the only fungal genus found to be significantly associated with yeast OTUs. *Amanita*, a speciose Basidiomycota genus containing many poisonous and edible mushroom species, was found to be associated with the Basidiomycota OTU *Vanrija humicola* (*P*_adj_= 0.03) and the Saccharomycotina *Te. cretensis/ kruisii* spp. complex (*P*_adj_= 0.035). Ten additional plant genera were associated with a single OTU each (*Vaccinium* – berry-producing shrubs, *Cu. cygneicollum, P*_adj_ = 0.035; *Thuja* - cypress trees, *D. hansenii, P*_adj_= 0.033; *Taxus* - yew trees, *Candida coipomoensis, P*_adj_< 0.0001; *Taraxacum* – dandelions, *Zygowilliopsis californica, P*_adj_< 0.001; *Rubus* – berry-producing bushes, *H. uvarum, P*_adj_= 0.048; *Pinus* – pine trees, *Schwanniomyces polymorphus, P*_adj_< 0.0001; *Morus* – mulberries, *Papiliotrema flavescens, P*_adj_= 0.035; *Festuca* – perennial tufted grasses, *Wickerhamomyces onychis, P*_adj_= *0*.015; *Betula* - birch trees, *Kwoniella betulae, P*_adj_= 0.018; *Alnus* - alder trees, *Nakazawaea anatomiae/populi* spp. complex, *P*_adj_< 0.0001).

Associations were examined at both the level of general substrate type and substrate genus because these categories do not overlap entirely. For example, the association of *L. thermotolerans* and the *Quercus* genus of oaks would have been missed at the substrate level, as *L. thermotolerans* had independent isolations from soil sixteen times, bark eight times, plant matter five times, as well as other various substrates like duff and leaves. Nonetheless we did find corroboration between the two analyses. This consistency was most obvious among the five OTUs that were associated with a biotic substrate and also with a corresponding genus into which that substrate falls: *Zygowilliopsis californica* (*Taraxacum* and flower), *Sy. polyspora* (*Picea* and needles), *L. kluyveri* (*Cercis* and bark), *H. uvarum* (*Rubis* and fruit) and *Te. cretensis/kruisii* spp. complex (*Amanita* and fungus). These analyses were performed independently on non-identical datasets (**Fig. S9A-B**), and so these corroborative findings lend high confidence in these associations. However, in the case of *H. uvarum*, this consistency is slightly complicated by additional associations. As mentioned, *H. uvarum* was associated with *Rubus* at the plant genus level and fruit at the substrate level, but we also found an additional substrate association of *H. uvarum* with fungus. Thus, while we observed corroboration of *H. uvarum*’s well-known role in rotting fruit fermentation, we also identified a potentially broader niche for this OTU (Albertin et al. 2016; Spencer, Spencer, De Figueroa and Heluane 1992).

In contrast to many previous studies (Naumov 1996; Naumov, Naumova and Sniegowski 1998; Sniegowski et al. 2002; Kowallik and Greig 2016; Wang et al. 2012; Zhang, Skelton, Garner and Goddard 2010), we did not find associations between *Saccharomyces* species and the oak genus *Quercus*. We isolated *S. cerevisiae* 21 independent times with only five isolations from *Quercus*-associated substrates (**Suppl. Table 9**). *S. paradoxus* was isolated 43 independent times with 7 isolations from *Quercus* substrates (**Suppl. Table 9**). The absence of associations cannot be explained by lack of sampling, as *Quercus* was the most deeply sampled plant genus in our dataset (**Fig. S8B**). The absence of statistical associations is noteworthy as oak trees have long been thought of as the natural habitat of *Saccharomyces* species, particularly *S. cerevisiae* and *S. paradoxus* (Kowallik and Greig 2016; Sniegowski et al. 2002; Wang et al. 2012; Zhang et al. 2010). Indeed, a previous study from our lab using a much smaller subset of these data did find a statistical association between the *Saccharomyces* and *Quercus* genera (Sylvester et al. 2015). This significant association was a result of direct hypothesis testing as opposed to the presently applied query of all possible associations. To ensure that these different results were due to increased sampling and not to differences in statistical methods, we applied the same statistics used to detect a *Quercus* - *Saccharomyces* association in Sylvester et al. 2015 and failed to recover a significant association in our larger dataset. Further, if Sylvester et al. 2015 had queried all possible associations using our permutation analysis, the association would have been non-significant. Because of the extensive literature focused on isolating *Saccharomyces* from oak trees, the assumption that oak exudate is a primary habitat of *Saccharomyces* species has percolated pervasively into the broader yeast literature. While it is obvious that *Saccharomyces* species can indeed be isolated from from oak trees and surrounding soils and leaf litter, it is also clear from our data that *S. cerevisiae, S. paradoxus*, and other *Saccharomyces* can be found on a wide array of substrates. The abundance of *S. cerevisiae* isolated outside oak-associated habitats has been noted before, leading to the suggestion that *S. cerevisiae* may be a widely distributed generalist, as opposed to occupying narrower sugar-rich niches, such a fruit and oak exudate (Goddard and Greig, 2015). Our findings provide further evidence that *S. cerevisiae*, and *Saccharomyces* spp. more broadly, occupy diverse habitats and are not confined to the bark of oak and related trees.

Studies wielding sufficient power to statistically associate yeast OTUs with habitat-types in a similar manner to our analyses are still rare, and therefore most of what is known about substrate preferences of yeasts come from a few isolations or anecdotal evidence. Still, many of the significant substrate associations we found in our data confirmed previously reported associations based on fewer isolations. A novel species previously described by our laboratory, *Kw. betulae*, was named for the birch genus from which it was isolated (Sylvester et al. 2015). The presently analyzed data, which contains the same isolates presented in the original description paper and several new isolates, found a statistical association with the *Betula* genus. *Sy. polyspora* is a known conifer pathogen, and we found it was statistically associated with the pine genus *Picea*, as well as with needles (Guertin, Zitouni, Targuay, Hogue and Beaulieu 2018; Talgø et al. 2010). The association we found between the opportunistic human pathogen *P. kudriavzevii* and fruit has been reported several times before (Douglass et al. 2018; Kurtzman, Fell and Boekout 2011; Opulente et al. 2019). Additional previously described associations that we replicated included *T. delbrueckii* and soil (Kurtzman, Fell and Boekhout 2011), the xylose-consuming yeast *Sc. ergatensis* and bark (Kurtzman, Fell and Boekhout 2011), *H. uvarum* and fruit (Hamby, Hernández, Boundy-Mills and Zalom 2012; Vadkertiová et al. 2012), and *Tr. porosum* and soil (Middelhoven, Scorzetti and Fell 2001). We also found logical indirect connections between the associations we found and previously reported isolation substrates. For example, *Su. bolitotheri* and *Te. cretensis/Te. kruisii* were originally isolated from the guts of basidiocarp-feeding beetles (Suh, McHugh and Blackwell 2004), and we found them both associated with fungi. *Scl. sphagnicola* is considered a moss pathogen (Tsuneda, Chen and Currah 2001), and we found it associated with lichens, an association that may be due to the similar habitats of mosses and lichens in the boreal forest from which it was isolated. Conversely, some associations found in these data were novel. *H. uvarum* has not been previously described as being a common yeast of macrofungi, but we found a robust association between the two. *Cu. cygneicollum* has been isolated from diverse substrates across the globe (Sampaio et al. 2004), but, to our knowledge, it has not been associated with fruit before.

#### Isolation temperature – higher taxonomic group associations

Isolations in this dataset were performed at one of four temperatures: 4°C, 10°C, 22°C, or 30°C. In total, isolation temperature was confidently assigned for 1,750 isolations. We have previously shown that isolation temperature dramatically affects taxonomic representation among isolates (Sylvester et al. 2015), and we found the same trends in the current data. Significant enrichment for Basidiomycota was observed at cooler isolation temperatures of 4°C (*P*_adj_= 1.75×10^−35^) and 10°C (*P*_adj_= 1.00×10^−54^, **Fig. 2A, Suppl. Table 10**). Similarly, an enrichment for Saccharomycotina was observed at 22°C (*P*_adj_= 2.45×10^−23^) and 30°C (*P*_adj_ = 8.90×10^−41^).

**Figure 2.**
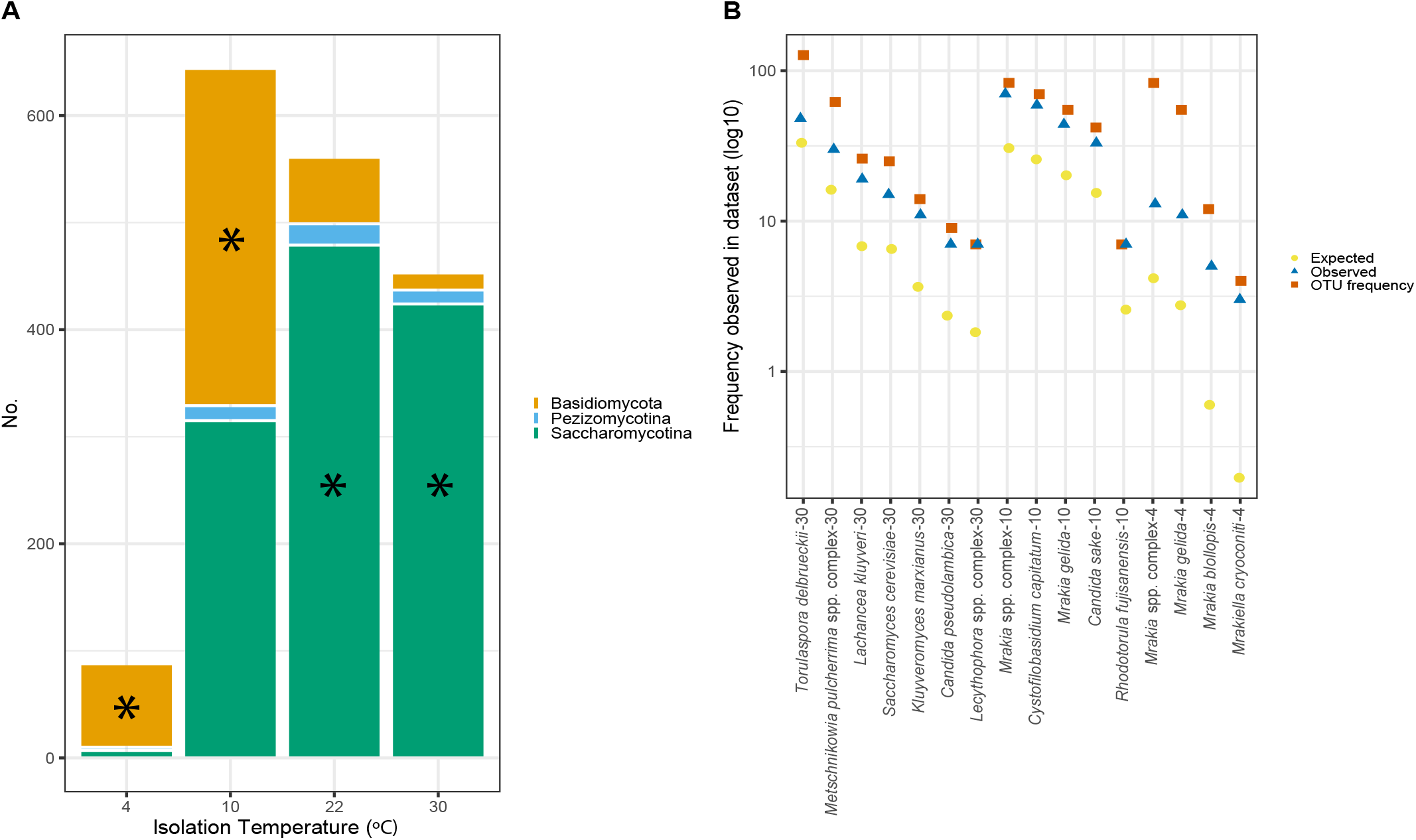
A) Distribution for the 1,750 isolations for which isolation temperature was known. Colors correspond to the taxonomic group of the corresponding isolate. * Basidiomycota OTUs were enriched at 4°C (*P*_adj_=1.75×10^−35^) and 10°C (*P*_adj_=1.00×10^−54^), and Saccharomycotina OTUs were enriched at 22°C (*P*_adj_=2.45×10^−23^) and 30°C(*P*_adj_ =8.90×10^−41^). **B)** Permutations identified 16 yeast OTU – isolation temperature associations observed more often than expected (*P*_adj_<0.05). Blue triangles indicate the number of times each combination was observed, red squares indicate the number of times the yeast OTU was observed in the permuted dataset, and yellow circles indicate the expected value of each association. The order of the results are by isolation temperature, followed by the number of isolations for each taxon-temperature combination.

#### Individual OTU – isolation temperature associations

To drill down on the specific yeast OTUs affected by isolation temperature, we examined our data for specific OTU – isolation temperature associations. Out of 444 yeast OTU – isolation temperature combinations (**Fig. S9C**), 16 significant positive associations were found (**Fig. 2B, Suppl. Table 11**). As expected, all OTUs associated with isolation at 4°C were Basidiomycota (*Mrakia blollopis, P*_adj_= 0.004; *Mrakiella cryoconite, P*_adj_= 0.014; *Mr. gelida, Mrakia* spp. complex, *P*_adj_< 0.0001). Five OTUs were associated with 10°C isolation, four of which were Basidiomycota (*Cystofilobasidium capitatum, Mr. gelida, Mrakia* spp. complex, *P*_adj_< 0.0001; *Rhodotorula fujisanensis, P*_adj_= 0.029), and one of which was a Saccharomycotina OTU (*C. sake, P*_adj_< 0.0001). There were no significant associations at 22°C. Seven total OTUs were found to be significantly associated with 30°C isolation, six of which were Saccharomycotina (*Kluyveromyces marxianus, L. kluyveri, P*_adj_< 0.0001; *M. pulcherrima* spp. complex, *P*_adj_= 0.008; *S. cerevisiae, P*_adj_= 0.014; *Candida pseudolambica, P*_adj_= 0.036; *T. delbrueckii, P*_adj_= 0.036). The remaining 30°C isolation–associated OTU was a dimorphic Pezizomycotina (*Lecythophora* sp., *P*_adj_= 0.004).

We have now repeatedly found taxonomic differences in isolation temperature, with Basidiomycota dominating cool isolation temperatures and Saccharomycotina dominating warm isolation temperatures (Sylvester et al. 2015). Here we found that taxonomic group effects emerge at the level of individual OTU – temperature associations as well. Eight of the nine significant associations with cold temperatures belonged to Basidiomycota. Conversely, five of the six yeasts associated with a warm isolation temperature belonged to Saccharomycotina and none to Basidiomycota. One significant exception to the higher taxonomic divisions among isolation temperatures was *C. sake*, a Saccharomycotina yeast significantly associated with 10°C isolations. *C. sake* is known as a cold-tolerant species and has been isolated from Arctic environments (Ballester-Tomás et al. 2017). Neither of the two known cold-tolerant *Saccharomyces* species in our dataset had significant isolation temperature associations, but both species were more successfully isolated at room temperature or slightly lower. *S. eubayanus* was isolated at about the same rate at 10°C (6 isolations) and 22°C (5 isolations) and never isolated at 30°C or 4°C, while *S. uvarum* was isolated only once at 10°C.

Germination, growth rates, and microbial competition are but a few of the processes likely to affect species-specific isolation success at different temperatures. The strong phylogenetic effects we found suggest that intrinsic biological properties drive most of the variation in isolation success. In confirmation of our power to detect biologically real temperature preferences, we recovered *S. cerevisiae* as a 30°C-associated OTU. These differences are useful for informing isolation approaches for different species and may point to differing ecology of these groups, as well as. For example, we do not recover the same warm-temperature association for *Saccharomyces paradoxus* that we do for *Saccharomyces cerevisiae*, and this corresponds well with the inferred different temperature-driven distributions of these two sister species (Robinson, Pinharanda and Bensasson 2016).

### Substrate and temperature diversity

#### Variation in diversity by substrate sampled

Maximizing taxonomic diversity recovered when sampling would be advantageous for yeast ecologists and taxonomists. To determine which substrates yield the highest within-substrate (alpha) diversity of yeasts, we estimated alpha diversity of each of the 15 substrates that were sampled at least 5 times using the Shannon-Wiener index (H’) (**Fig. 3A, Suppl. Table 12**). We corroborated diversity estimates by inspecting rarefaction curves by substrate (**Fig. 3B-C)**. Given the effect of sampling density on H’, the most useful comparisons are among substrates with similar sampling densities. As expected, the substrates with the highest H’ were bark and soil, the most densely sampled substrates. Nonetheless, bark exhibits a slightly higher H’ index than soil, despite having ≈100 fewer processed samples. Higher diversity on bark relative to soil may be an ecological property unique to yeasts among fungi, as a study examining metagenome datasets recently found soil to have higher total fungal diversity than shoots or wood (Baldrian, Vĕtrovský, Lepinay and Kohout 2021). Fungi, duff, plant matter, fruit, and leaves were all sampled to similar extents, but fungi yielded lower diversity than similarly sampled substrates, while leaves showed a relatively high diversity. Amongst the most sparsely sampled substrates, pine cones had elevated diversity relative to nuts, wood, needles, and twigs.

**Figure 3.**
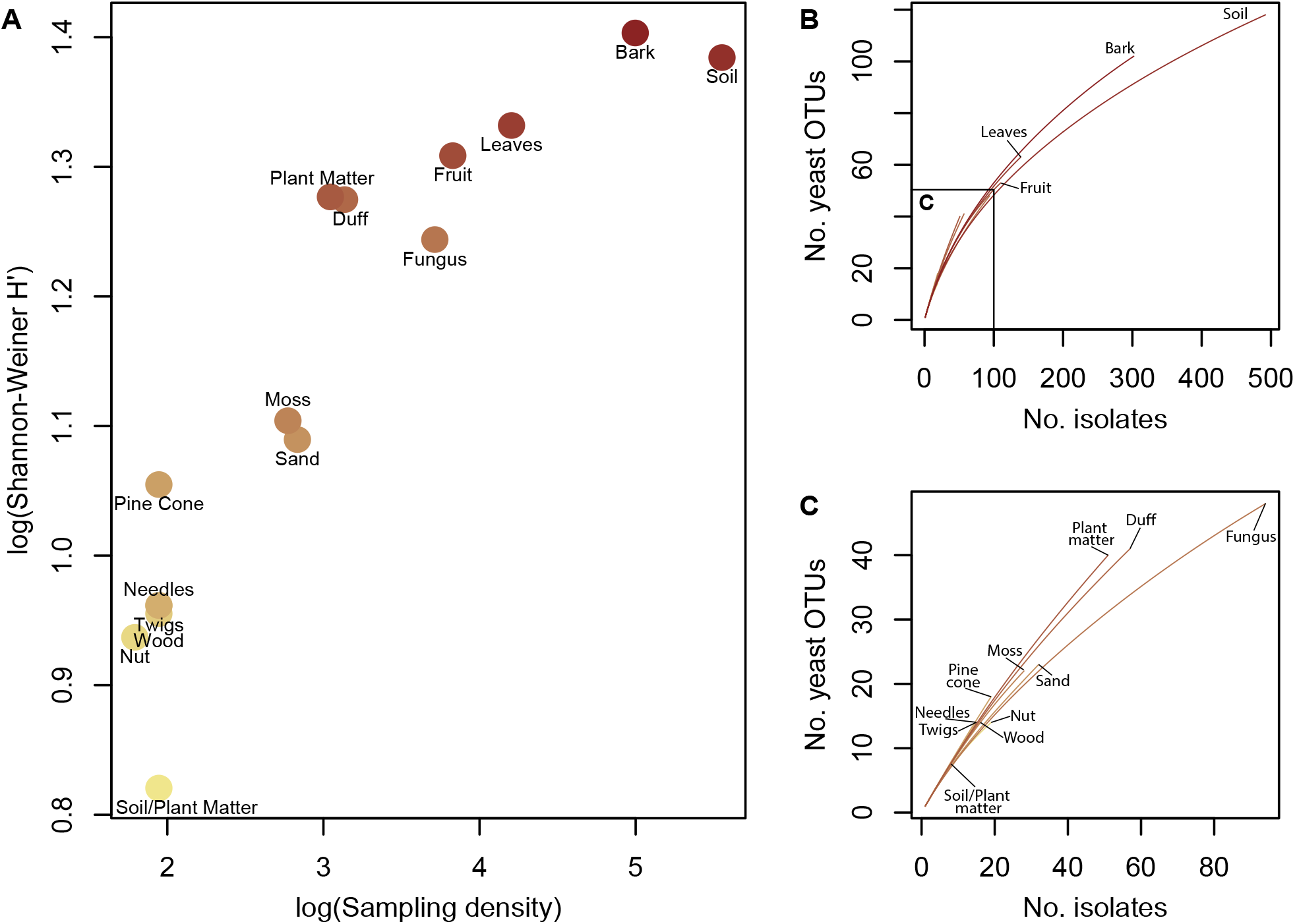
A) Shannon-Wiener (H’) indices were calculated for substrate categories sampled at least five times. Points are labeled by substrate type, and the color gradient of the points corresponds to increasing diversity. Sampling density affects the H’ index estimate, but some substrate categories have similar sampling densities with very different H’ indices. **B)** Rarefaction curves for each substrate category sampled at least five times; most distinguishable are bark, soil, leaves, and fruit. **C)** The same rarefaction curves as shown in panel B are zoomed in to show substrates with fewer isolates.

#### Variation in diversity by isolation temperature

Diversity isolated from samples may also depend on enrichment temperatures, so we used a similar approach to examine the alpha diversity among each enrichment temperature. We found 22°C to have the highest level of diversity with an index value of 4.3 and 4°C to have the lowest level of diversity with an index value of 3.21 (**Fig. 4A&C, Suppl. Table 13**). Because isolation temperature had a drastic effect on the subphyla of resultant isolates, we used the same approach to analyze diversity individually within each taxonomic group (**Fig. 4D-F, Suppl. Table 13)**. Among OTUs within Saccharomycotina, H’ indices were similarly high at 30°C, 22°C, and 10°C, but dropped off sharply at 4°C. The reciprocal pattern was seen in Basidiomycota, which exhibited a sharp drop off at 30°C. Pezizomycotina showed overall low diversity with an optimum at 22°C. The low diversity here likely reflects our protocol’s intentional avoidance of mold-like colonies, a morphology common in Pezizomycotina. Even when considering higher taxonomic units separately, maximum diversity was always achieved at 22°C. This result, coupled with the finding that no individual taxonomic unit was preferentially isolated at 22°C, suggests that intermediate temperatures are ideal for generally querying the yeast diversity present in a given sample.

**Figure 4.**
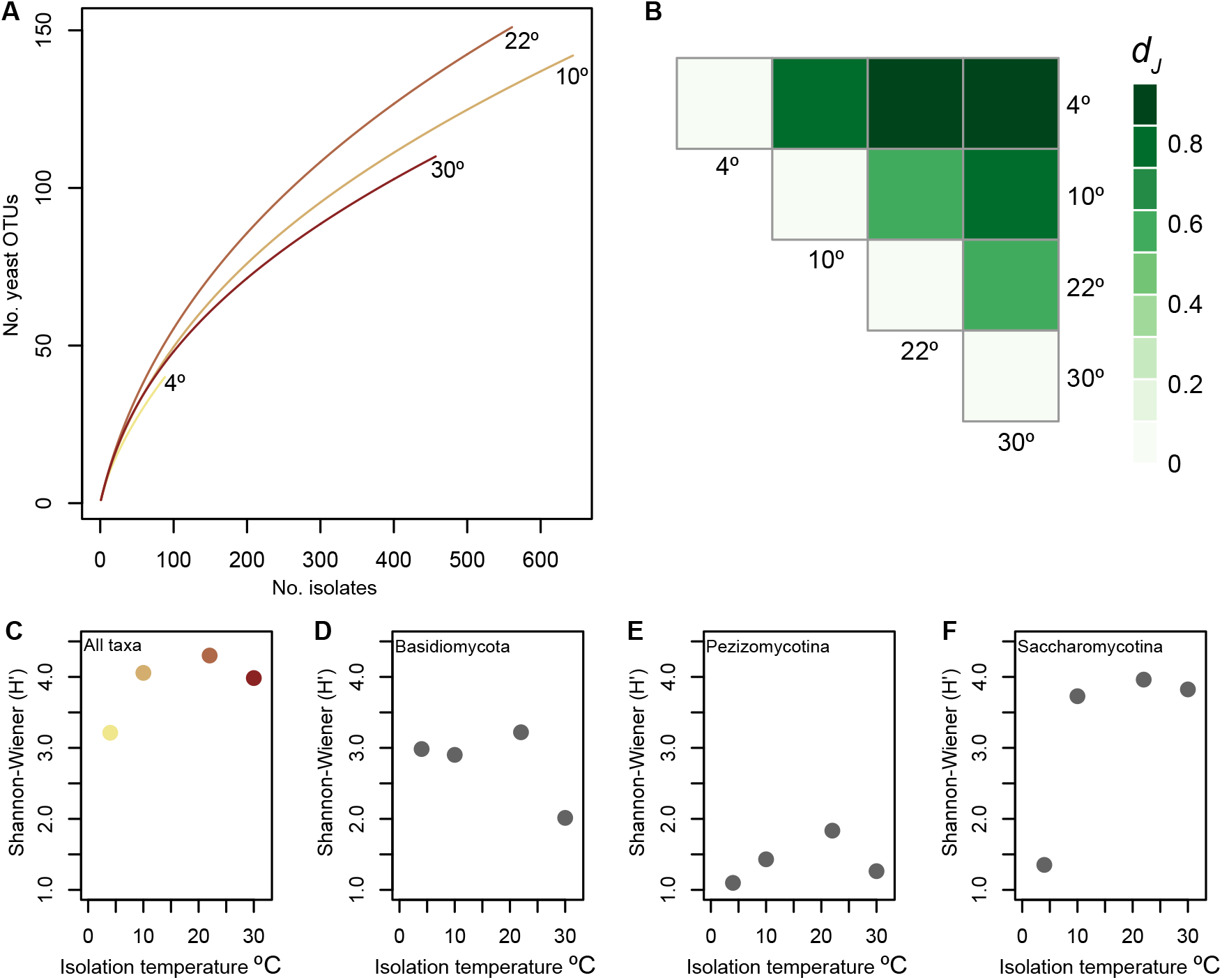
A) Rarefaction curves of each isolation temperature. The color gradient corresponds to increasing temperature. **B**) Heatmap of Jaccard distance (*D*_*J*_) between isolation temperatures. **C**) Shannon-Wiener (H’) indices of each isolation temperature for all taxa. The color gradient corresponds to increasing temperature. Isolations at 22°C are optimal for maximizing yeast diversity. 4°C isolations have reduced diversity. **D-F)** Shannon-Wiener (H’) indices of higher taxonomic ranks: Basidiomycota (**D**), Pezizomycotina (**E**), and Saccharomycotina (**F**). Basidiomycete diversity was maximized at lower temperatures, while Saccharomycotina diversity was maximized at higher temperatures.

Both the yeast taxon-isolation temperature associations and the alpha diversity analyses reported above are indirect evidence that different isolation temperatures yield very different OTU compositions. To directly test this hypothesis, we examined beta diversity amongst isolation temperatures using Jaccard distance (*D*_*J*_) (**Fig. 4B, Suppl. Table 14)**. As expected, there was very little overlap between the assemblages of taxa recovered from isolations at 4°C and 30°C (*D*_*J*_= 0.95). *D*_*J*_ values overall correlated roughly, but not precisely, with the difference between isolation temperatures. For example, *D*_*J*_ between 10°C and 4°C (0.84) was higher than between 22°C and 30°C (0.63). Therefore a drop from 10°C to 4°C isolations had a greater effect on the taxa recovered than an increase from 22°C to 30°C, despite the former having a difference of 6° and the latter having a difference of 8°. The lowest beta diversity was observed between 22°C and 10°C, indicating that these isolation temperatures impose a similar thermal selection. The low beta diversity observed between 22°C and other isolation temperatures (excluding 4°C) further supports the use of isolation temperatures near 22°C to maximize the recovery of yeast taxonomic diversity.

### Conclusions: A window into yeast ecology reveals geospatial variation, habitat specificity, and diverse thermal niches

Inferences regarding the distributions, habitat preferences, and broader ecological roles of yeasts have proven difficult to generate due to the low abundance of yeasts in nature and the tedium associated with detecting them. Here, we leveraged a dataset of nearly 2,000 natural yeast isolates, the largest such collection accrued using the same standardized isolation protocols and conditions, to gain insight into patterns of yeast distribution, habitat specificity, and temperature associations. While many yeast taxa were repeatedly isolated in our dataset, we found that only 11 such taxa were always isolated in regions where sampling should have been dense enough to detect them. Our analysis does not exclude any OTU from being cosmopolitan, rather we present a simple method for objectively detecting those taxa that are broadly distributed across regions. We similarly employed an objective statistical method of detecting associations between yeast OTUs and sampled substrates and found some yeast OTUs to exhibit strong patterns of substrate specificity. When the same approach was applied to isolation temperature, we found a significant effect of incubation temperature on the phylogenetic groups ultimately isolated, a result we had reported before (Sylvester et al. 2015) that is firmly reinforced here. Finally, we examined the overall yeast diversity achieved from different substrates and using different incubation temperatures, and found bark samples and room temperature incubations to yield the highest yeast diversity. Taken together, these results paint a picture of ecological structure in the distribution, habitat, and temperature preferences of yeasts.

All data is provided in the form of supplemental tables.

Suppl. Table 1 – Raw metadata for 1,962 isolates.

Suppl. Table 2 – OTUs isolated only once.

Suppl. Table 3 – OTUs isolated greater than 20 times.

Suppl. Table 4 – Discovery rates and regional expectations for 27 OTUs.

Suppl. Table 5 – Cosmopolitan OTUs.

Suppl. Table 6 – Contingency table for higher taxonomic group representation across substrate types.

Suppl. Table 7 – Permutation results for specific substrate – yeast OTU associations.

Suppl. Table 8 – Permutation results for substrate genus – yeast OTU associations.

Suppl. Table 9 – All isolates of *Saccharomyces cerevisiae* and *Saccharomyces paradoxus*.

Suppl. Table 10 – Contingency table for higher taxonomic group representation across isolation temperatures.

Suppl. Table 11 – Permutation results for isolation temperature – yeast OTU associations.

Suppl. Table 12 – Shannon-Weiner indices for 15 substrate types.

Suppl. Table 13 – Shannon-Weiner indices for 4 isolation temperatures.

Suppl. Table 14 - Jaccard distance matrix of isolation temperatures.

## Acknowledgments

We thank all past and present participants in the Wild YEAST Program (http://go.wisc.edu), including all individuals that contributed samples to this study: Alison Coffey, Alysha Heimberg, Anita R. & S. Todd Hittinger, Anna S. Kropornicka, Kurt Ehlert, Ashanti Rogers, Beth Sargent, Bill Saucier, Bill Vagt, Brian P. H. Metzger, Meihua Christina Kuang, D. Leith Nye, Drew T. Doering, EmilyClare P. Baker, Erik D. Jessen, Erica L. Macke, Harry Iwatsuki, Jacob Barnes, Jin Kang, John E. Pool, Katie Burjek, Laura Beck, Leslie Shown, Mary B. O’Neill, Russell L. Wrobel, William G. Alexander, S. Smead, Mike Siegel, the Zasadil family, and Mel Langdon. This material is based upon work supported by the National Science Foundation under Grant Nos. DEB-1253634 (to C.T.H.), DEB-1442148 (to C.T.H.), and DGE-1256259 (to Q.K.L); in part by the DOE Great Lakes Bioenergy Research Center (DOE BER Office of Science DE-SC0018409); and the USDA National Institute of Food and Agriculture (Hatch Project 1020204). W.J.S. was supported by a University of Wisconsin- Madison Genetics and Genomics Undergraduate Distinguished Research Fellowship while completing this work. K.J.F. is a Morgridge Metabolism Interdisciplinary Fellow of the Morgridge Institute for Research. Q.K.L. was also supported by the Predoctoral Training Program in Genetics, funded by the National Institutes of Health (Grant No. 5T32GM007133). C.T.H. is a Pew Scholar in the Biomedical Sciences and an H. I. Romnes Faculty Fellow, supported by the Pew Charitable Trusts and Office of the Vice Chancellor for Research and Graduate Education with funding from the Wisconsin Alumni Research Foundation, respectively.

## Figure Legends

**Figure S1**) Histogram of isolates in the complete dataset (top). When the 1,962 isolations were filtered to remove duplicate OTUs derived from the same processed sample, 1,518 unique isolations remained (bottom).

**Figure S2**) The distribution of isolations in the dataset by climate region. The number of unique isolations (upper, bold) and the number of unique taxonomic units (lower, italics) are shown. Climate regions correspond to the 9 climatically consistent regions of the contiguous US identified by NOAA (https://www.ncdc.noaa.gov/monitoring-references/maps/us-climate-regions.php). We included the additional category of Arctic for Alaskan samples.

**Figure S3) A)** Rarefaction curve for 688 unique samples (black line) with 95% confidence intervals. **B)** Rarefaction of 688 unique samples by the higher taxonomic units Basidiomycota (yellow), Pezizomycotina (blue), and Saccharomycotina (green).

**Figure S4)** Taxonomic representation across unique isolations (light grey, top count) and unique OTUs (dark grey, bottom count). The dearth of Pezizomycotina OTUs is likely reflective of our isolation protocol, in which the selection of mold-like colonies is deliberately avoided.

**Figure S5)**116 singleton OTUs were isolated just once. **A)** Distribution of substrate categories among singletons. Singleton OTUs were not enriched for any substrate type. **B)** Singletons were enriched for OTUs belonging to the Basidiomycota (Padj=0.003) and Pezizomycotina (Padj=0.0005). **C)** Isolation locations of singletons.

**Figure S6)A**) The number of independent isolations of each cosmopolitan OTU. **B**) The number of climatic regions each OTU was detected in.

**Figure S7)** Cosmopolitan OTUs were defined as those OTUs that were isolated from regions where they were expected based on sampling density (allowing for one missed region). Eleven cosmopolitan OTUs were identified using this approach. **A)** Cosmopolitan yeasts were enriched in soil samples (*p*= 3.09e-05). **B)** Cosmopolitan yeasts all belong to Saccharomycotina or Basidiomycota, but there was no enrichment for taxonomic groups among cosmopolitan OTUs. **C)** Map of cosmopolitan isolation locations. OTUs are differentiated by color.

**Figure S8)** Forty discrete substrate categories were annotated for 1,522 isolations. **A)** Distribution of unique isolations among substrate categories. Category sampling was uneven. **B)** Substrate genera could be assigned to 1,026 isolations of diverse substrate categories. For each genus, the type of substrate collected is shown with different color bars. Substrate categories were either directly harvested from the substrate (e.g. pine needles from pine tree) or indirectly associated with the substrate (e.g. soil from the base of a pine tree).

**Figure S9)** Top) Observed data fed into substrate association permutations. Middle) Observed data fed into substrate plant genus association permutations. Bottom) Observed data fed into isolation temperature 0 associations.

## Notes

### Competing Interest Statement

The authors have declared no competing interest.

