## Supplementary figures and images for "Substrate, temperature, and geographical patterns among nearly 2,000 natural yeast isolates"

### Supplementa Figure 1

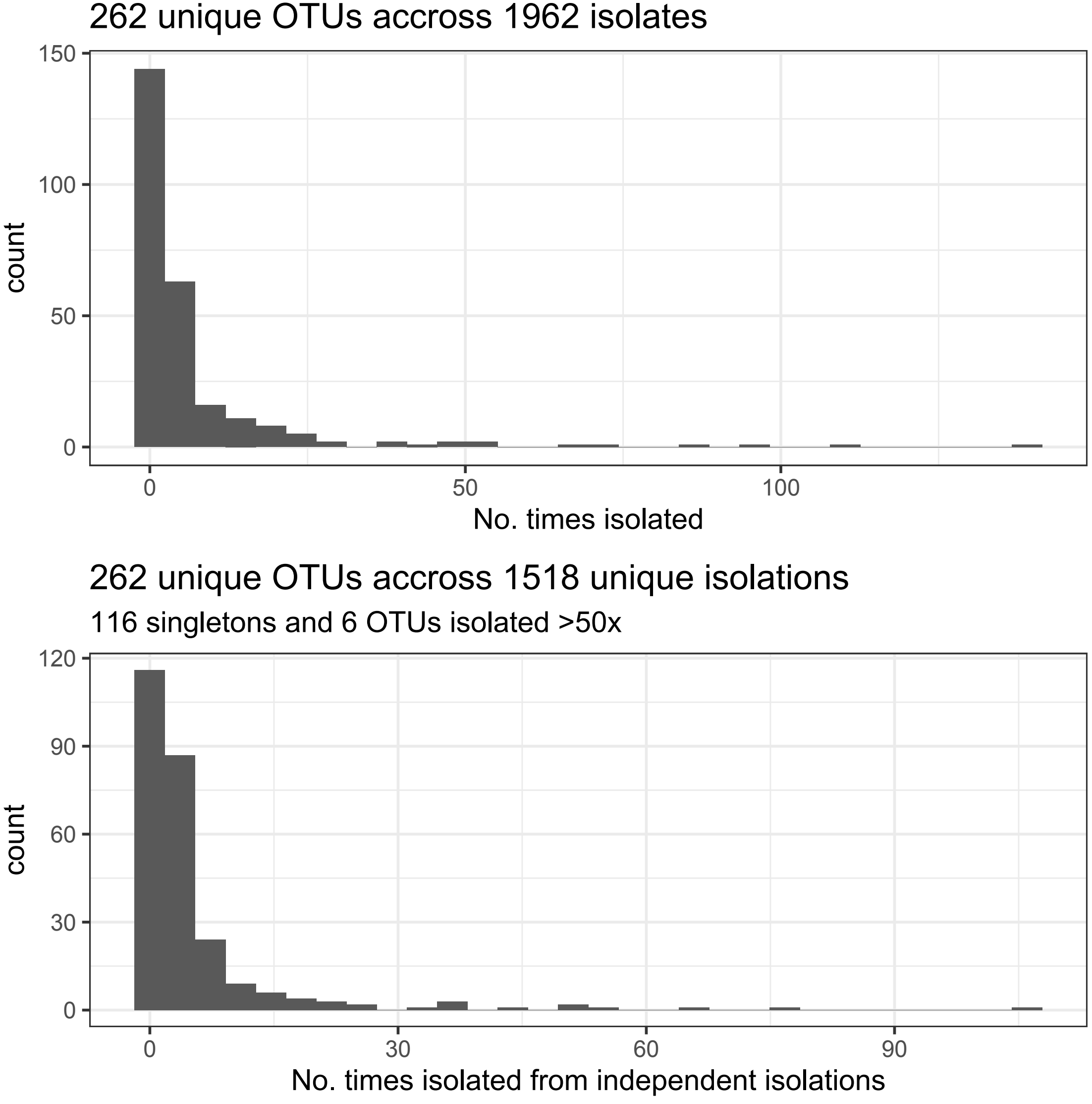

### Supplementa Figure 2

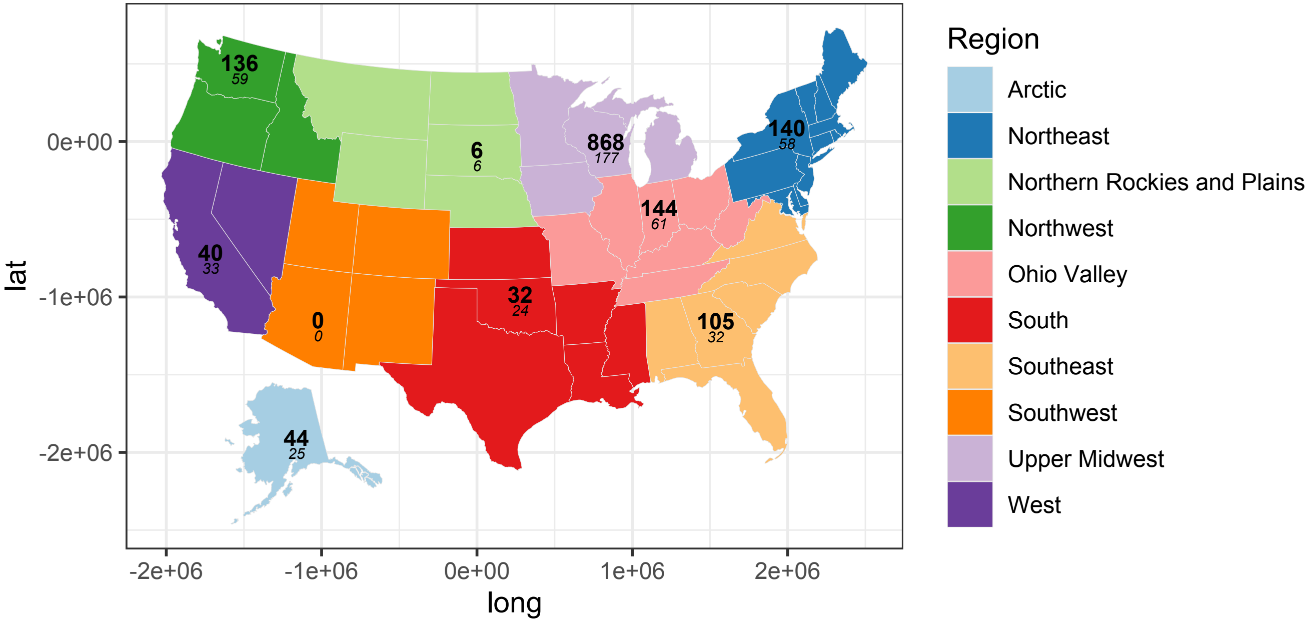

### Supplementa Figure 3

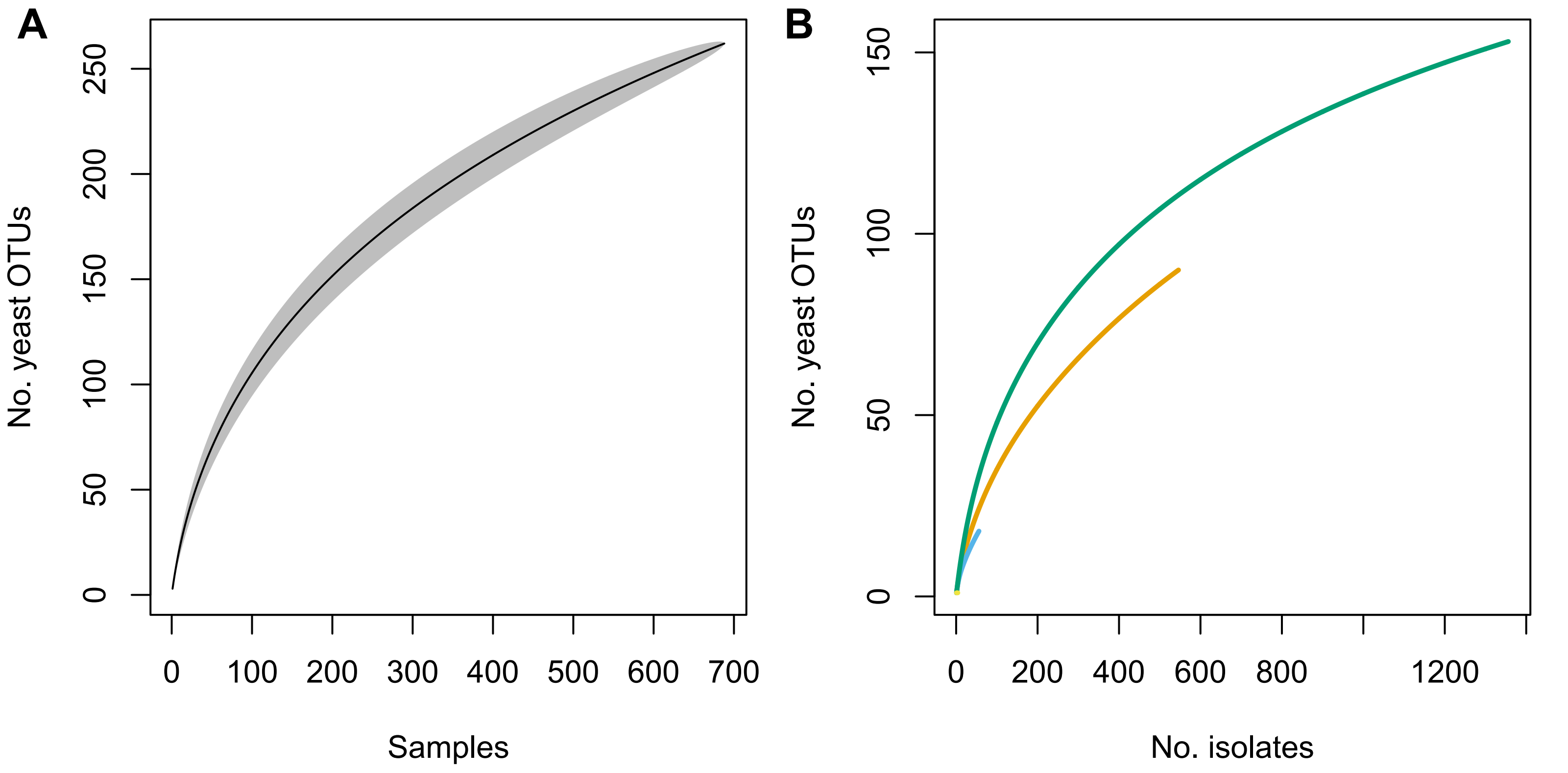

### Supplementa Figure 4

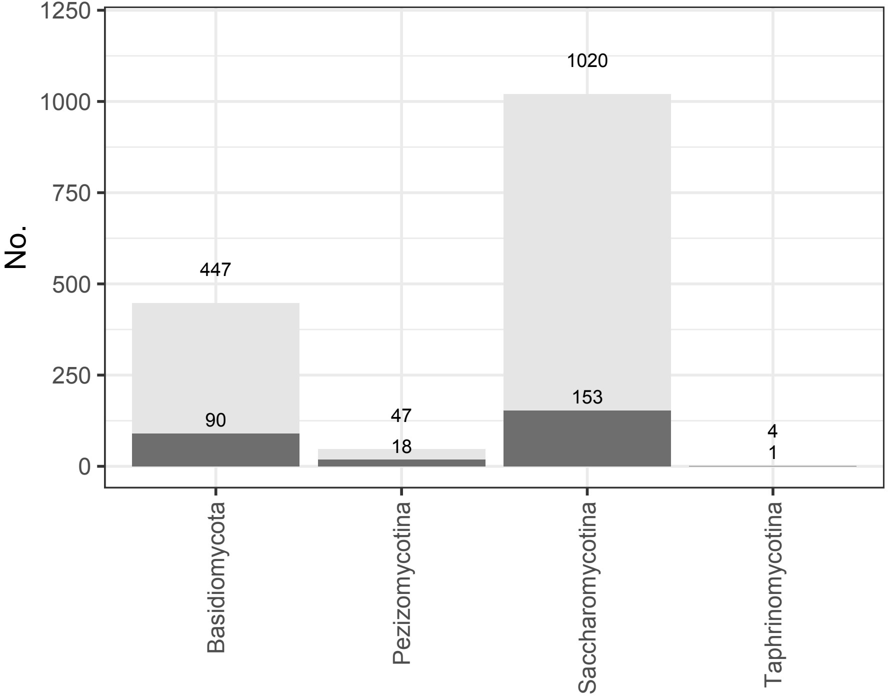

### Supplementa Figure 5

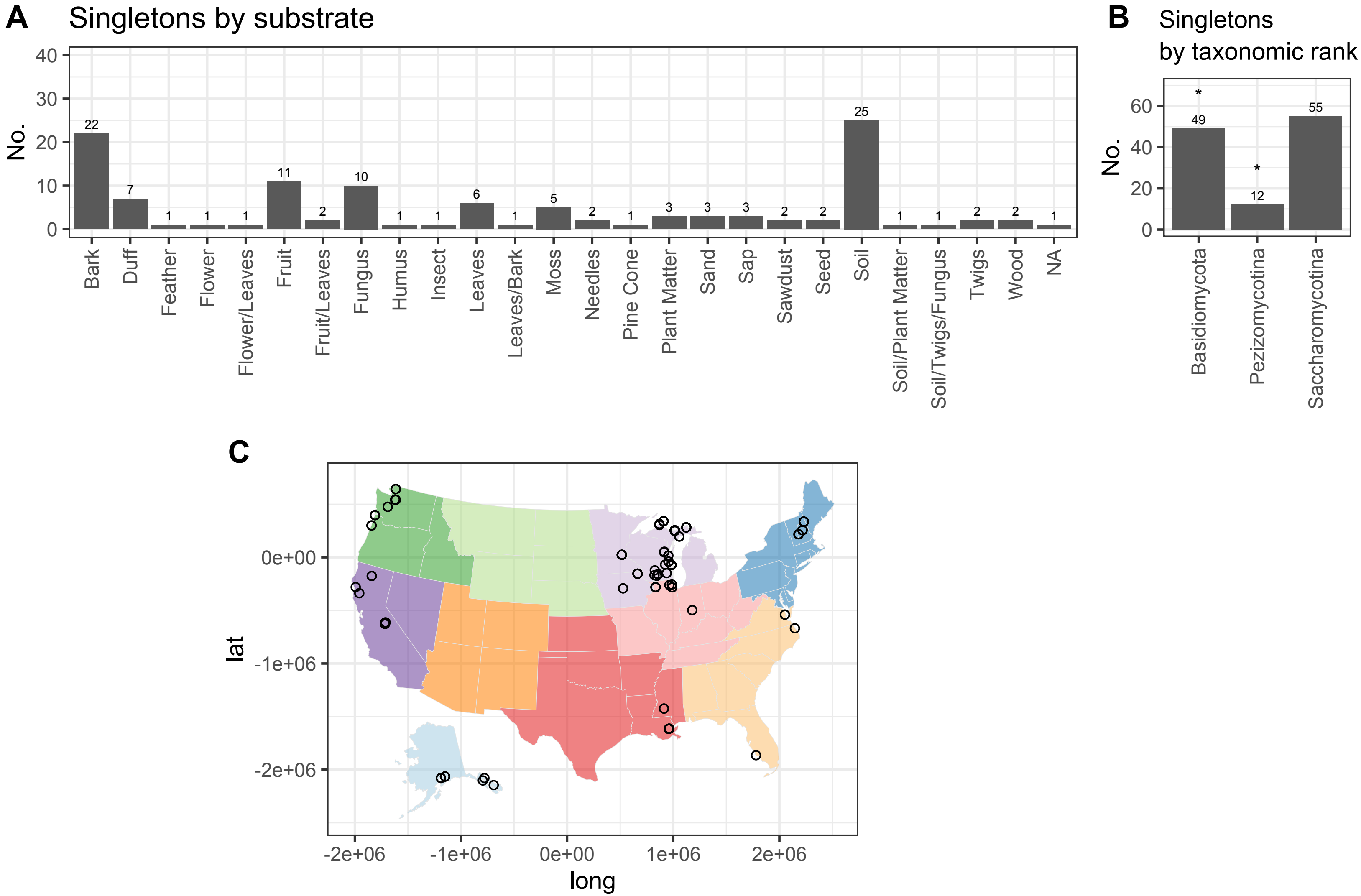

### Supplementa Figure 6

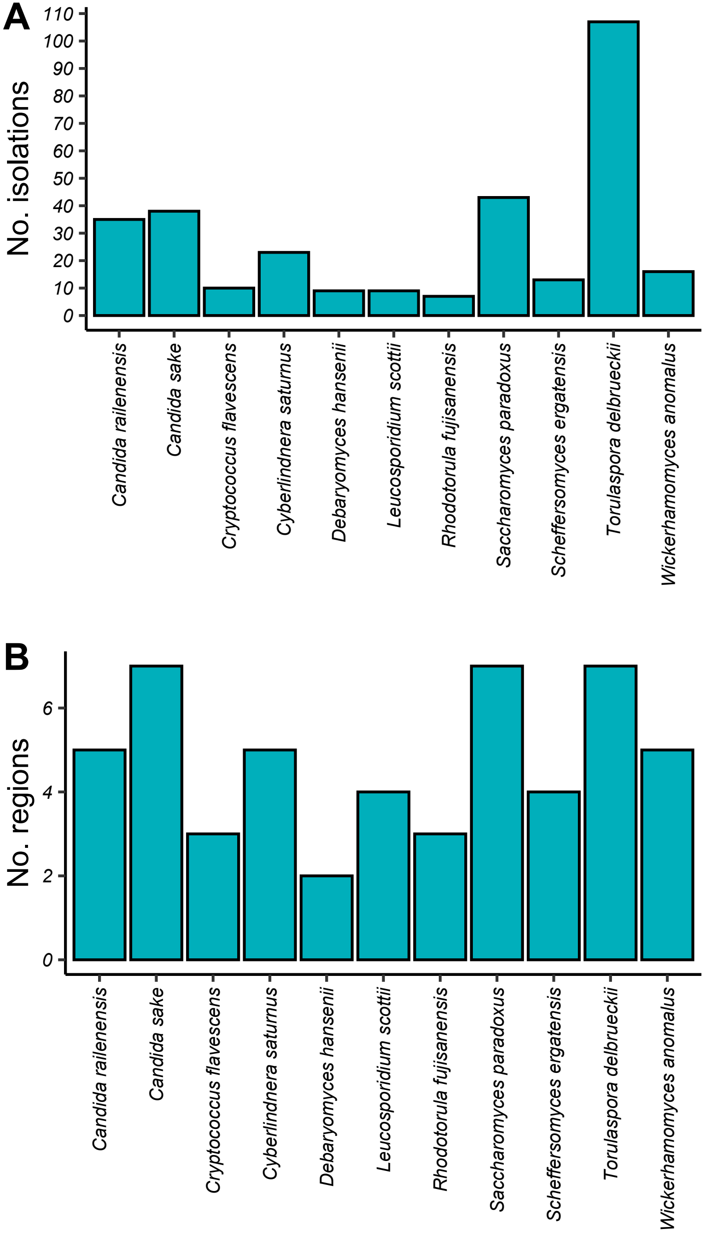

### Supplementa Figure 7

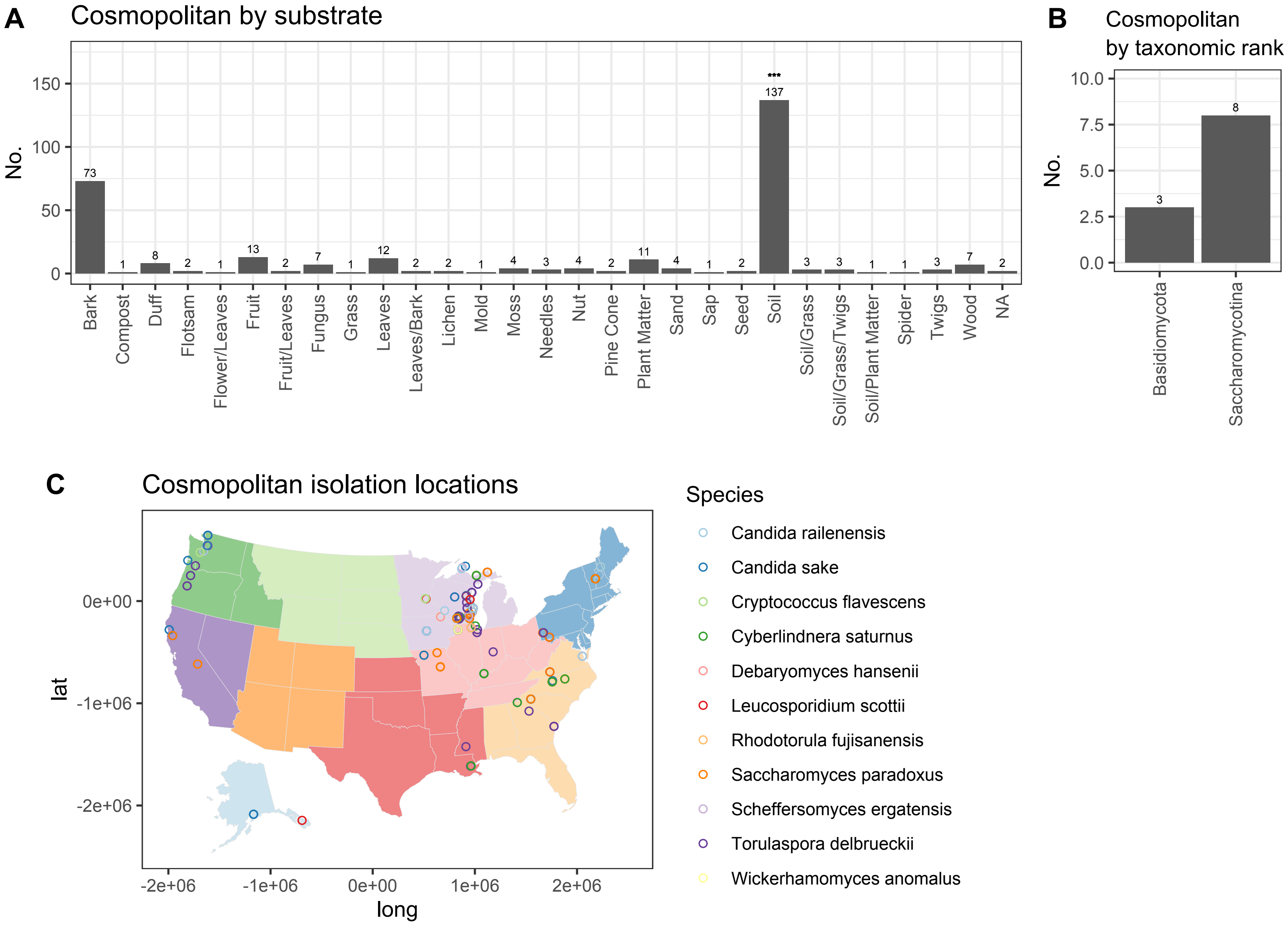

### Supplementa Figure 8

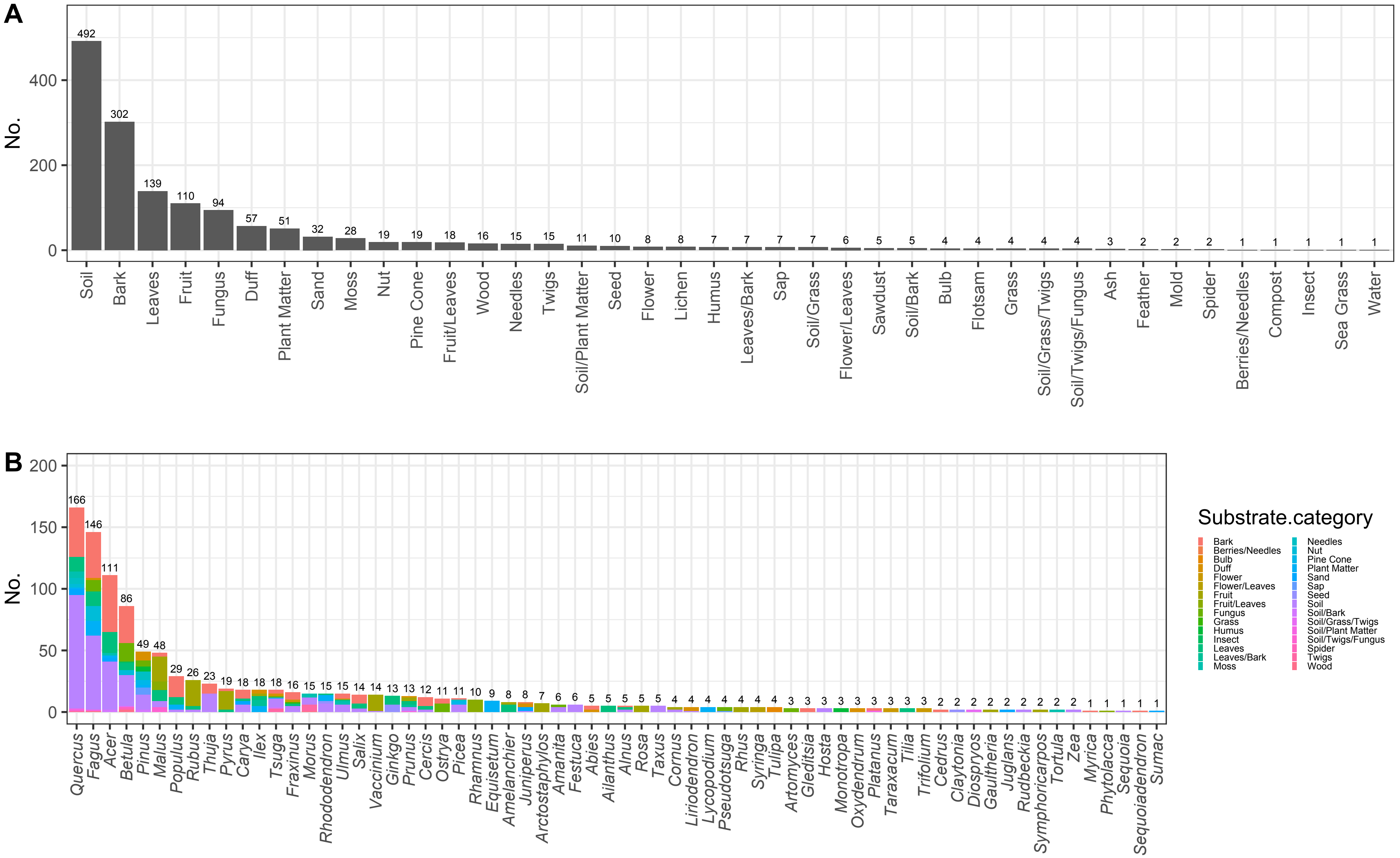

### Supplementa Figure 9

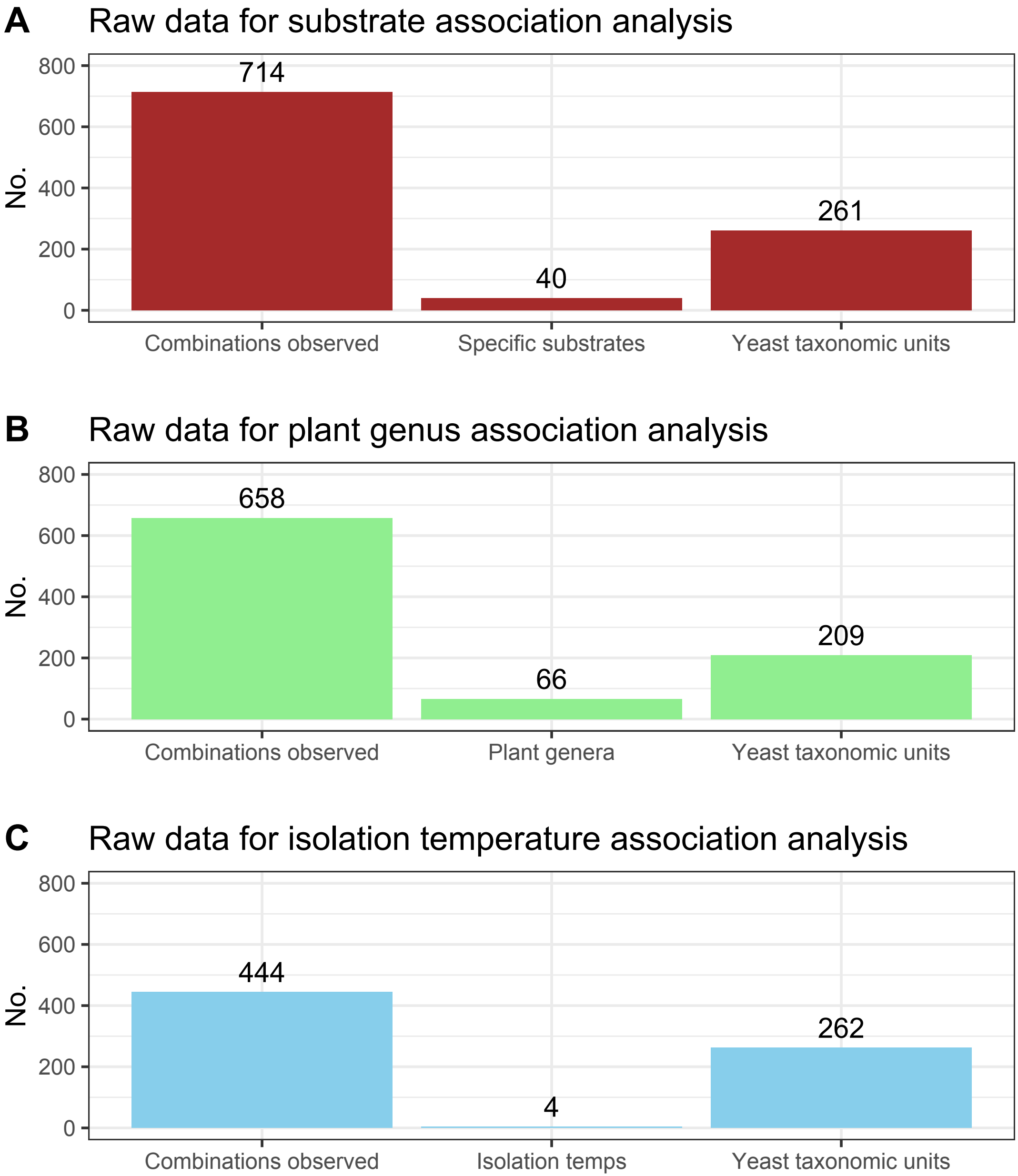
