## Supplementa Note 1 for "Substrate, temperature, and geographical patterns among nearly 2,000 natural yeast isolates"

**Supplementary Note 1**

**Yeast Enrichment, Culturing, Identification Protocols (Spurley/Fisher/Langdon...Opulente/Hittinger et al. (submitted to *Yeast*))**

*Modified from Sylvester et al. (2015) FEMS Yeast Res.*

**Collect sample**

Materials

- 1 sterile bag or tube per collected sample

1. Collect approximately 1 tablespoon of substrate in sterile bag or tube (without touching the sample with your hands).
2. Record sampling information –
   - Date, Location w/ GPS, Associated species (plant or fungi) if known.

**Inoculate into enrichment media**

Materials

- 3 15mL tubes/sample
- 3 negative control 15mL tubes
- enrichment media
  - ml enrichment media needed = ((3*# samples) + 3) x 9
- scoopula/tweezers

1. Label 3 15mL tubes for each sample (1 for each isolation temperature). Label 1 negative control tube for each isolation temperature.
   a. Sample #
   b. Initials
   c. Date
   d. Temperature (10°C, Room, 30°C)
   e. Sugar percentage
2. Near a flame, pipette 9mL enrichment media into each tube.
3. Use scoopula or forceps to load samples into tubes. Ethanol and flame sterilize scoopula and/or forceps between samples.
4. Vortex each tube for 3 seconds. Place in respective temperature.
5. Check regularly for growth - bubbles and white/whitish sediment.

**Passage cultures for second enrichment**

Materials

- 1 5mL tube (autoclaved) per sample/control
- enrichment media
  - ml enrichment needed = # tubes*4

1. Label autoclaved 5mL tubes with sample information, sugar percentage, date, and temperature.
2. Add 4mL of enrichment media to labeled tubes.
3. Vortex original 15 ml sample tubes with growth.
4. Pipette 10μL of liquid from 15ml-processed tube into the correspondingly labeled 5mL tube.
5. Place in respective temperature.
6. Watch for signs of growth - bubbles and white/whitish sediment.

**Plate enriched cultures for single colonies**

Materials

- two 1.5mL tubes per sample
- 1 YPD plate per sample
- sterile glass beads

1. Label two 1.5mL tubes for serial dilution for each sample.
   a. Tube 1 – sample # 10^-2^
   b. Tube 2 – sample # 10^-4^
2. Add 990 μL sterile H_2_O to each tube.
3. Vortex 5mL sample tubes and add 10μL to the first dilution tube. (10^-2^ or 1:100 dilution).
4. Vortex first dilution tube and add 10μL to the second dilution tube. (10^-4^ or 1:10,000 final dilution).
5. Label the outer edge of the bottom of YPD plates with tube information (initials, sample#, date, temperature, sugar percentage of enrichment media).
6. Pour ~10-15 sterile glass beads into the lid of the labeled plate.
7. Flip the plate with the lid on.
8. Lift the lid and pipette 100μL of vortexed 10^-4^ diluted suspension onto the glass beads. Replace lid and spread sample around the plate by shaking plate side to side.
9. Incubate plates (agar side up) in corresponding temperature.

**Streak to pure colonies of each morphotype**

Materials

- YPD plates
- toothpicks

1. Identify how many different morphotypes (different looking colonies) there are on each plate.
2. Choose one colony of each morphotype. Label each morphotype with a different letter starting with A.
3. Split a YPD plates into wedges (at least 4) for each distinct morphotype and label with date, temp, sample #, and morphotype letter.
4. Use a toothpick/stick to streak out a representative of each morphotype isolated.
5. Incubate upside down at respective temperatures (unless drippy then place upright).
6. Watch for single colony growth.
   *Note*: If your streaks overgrow and no longer have single colonies restreak from those cells to a new wedge.

**Inoculate culture for cryopreservation and DNA extraction**

Materials

- 1 glass culture tube per sample
- 3 mL YPD per sample
- toothpicks

1. Label 1 glass culture tube for each morphotype with sample info.
2. Dispense 3ml of YPD into each tube. Dispense a negative control tube as well.
3. Use a toothpick to pick a single colony. If there are no singles you must restreak. Smear the cells onto the inside side of the culture tube about two inches down the tube.
4. Vortex the tube so that the cell smear is washed into the media.
5. Balance on the culture wheel at the appropriate temperature.

**Freeze Down**

Materials

- 150 µL 50% glycerol per sample
- 1 cryopreservation tube per sample

1. Label each cryotube.
2. Add 150 μL of 50% glycerol.
3. Add 350 μL of saturated culture and mix by pipetting up and down.

*Note*: Vortex flocculating cultures vigorously to homogenize.

4. Put in micronics rack in -80°C freezer

**NaOH DNA extraction**

Materials

- 50 µL of 10mM NaOH per sample
- 0.2 mL strip tubes

1. Add ~200 μL of each saturated culture to the correct labeled strip tube.
2. Spin in picofuge to pellet.
3. Discard supernatant.
4. Add 50 μL of 10mM NaOH and pipette up and down or vortex to mix.
5. Include one negative control by adding NaOH to a tube with no cells in it
6. Incubate on thermocycler at 99°C for 15 min.
7. Store at 4°C.

**ITS amplification**

Materials

- 5X Taq Buffer (New England Biolabs)
- 10 mM dNTPs
- 10 μM ITS1 oligo stock
- 10 μM NL4 oligo stock
- Taq Polymerase (New England Biolabs)
- 0.2 mL PCR strip tubes

oligos:

ITS1 - TCCGTAGGTGAACCTGCGG

NL4 - GGTCCGTGTTTCAAGACGG

1. Prepare Master Mix on ice

| **Reagent** | **Volume per reaction** |
| --- | --- |
| H_2_O | 14.6 μL |
| 10x Taq Buffer | 2 μL |
| dNTPs | 0.6 μL |
| F oligo (ITS1) | 0.8 μL |
| R oligo (NL4) | 0.8 μL |
| Taq | 0.2 μL |
| NaOH DNA prep | 1 μL |

2. Mix well by pipetting up and down
3. Add 19μL of Master Mix to each sample’s strip tube, including the negative control (just boiled NaOH) tube.
4. Add 1μL NaOH prepped DNA for each sample, avoiding pelleted cells at the bottom of the tube.
5. Amplify on thermocycler.

Cycler conditions

94°C / 2 min

94°C / 2 min

40X	48°C / 30 sec

65°C / 3 min

65°C / 10 min

6. Verify a correct amplicon of ~1.4 kb through gel fractionation.

**Cleanup PCR**

Materials

- 10X Antarctic Phosphatase Buffer (New England Biolabs)
- Antarctic Phosphatase (New England Biolabs, 5,000 U/ml)
- Exonuclease I (New England Biolabs, 20,000 U/ml)
- 0.2 ml strip tubes

1. Prepare Master Mix on ice.

| **Reagent** | **Volume per reaction** |
| --- | --- |
| H_2_O | 1.8 µL |
| 10X Antarctic Phosphatase Buffer | 2.0 µL |
| Antarctic Phosphatase | 1.0 µL |
| Exonuclease I | 0.2 µL |

2. Add 5 µL of Master Mix to the remaining 15 µL of PCR reaction for each sample.
3. Incubate reactions at 37°C for 30 minutes.
4. Deactivate enzymes at 80°C for 20 minutes.
5. Store at 4°C.

**Sanger sequencing reaction**

Materials

- 5X BigDye Buffer (Applied Biosystems)
- BigDye reagent (Applied Biosystems)
- 10 μM ITS4 oligo

Oligos:

ITS4 – GCATATCAATAAGCGGAGGA

1. Make Master Mix on ice.

| **Reagent** | **Volume per reaction** |
| --- | --- |
| H_2_O | 5.75 µL |
| BigDye Buffer | 2.25 µL |
| BigDye | 0.4 µL |
| ITS4 primer | 0.6 µL |

2. Mix master mix well and distribute 9µL to each tube.
3. Add 1 µL of each Exo/AP product and mix by pipetting up and down.
4. Perform Sanger reaction on thermocycler.

Cycler conditions

94°C / 2 min

94°C / 10 sec

35X	52°C / 15 sec

60°C / 3 min

72°C / 1 min

**Bead**

Materials

- 250 µL 85% ethanol per sample
- Axygen DyeClean beads
- magnetic tube rack or plate

1. Prepare 85% ethanol fresh for each cleanup.
2. Thoroughly vortex to resuspend Axygen DyeClean beads
3. Make a bead-EtOH master mix:

| **Reagent** | **Volume per sample** |
| --- | --- |
| Axygen DyeClean beads | 5 µL |
| 85% EtOH | 42 µL |

4. Add 47μL of mixture from step 3 to each BigDye product. Pipette up and down to mix.
5. Place samples on magnetic plate for 5 min until beads separate from solution.
6. Remove the clear supernatant and discard.
7. Pipette 100μL of 85% ethanol over beads then remove.
8. Repeat 85% ethanol wash once more.
9. Remove samples from magnet.
10. Add 20 μL of ddH_2_O or TE buffer to beads and mix by pipette to resuspend beads.
11. Place samples back onto magnetic plate to separate beads.
12. Remove 20µL of the supernatant and transfer to a tube or plate for sequencing.

**Media Recipes**

**2X Synthetic Complete** (500 mL) – autoclave to sterilize

5.00g	Ammonium Sulfate

1.72g	Yeast Nitrogen Base w/o AA (YNB)

2.00g	Complete Drop-out mix

500 mL	milliQ H_2_O

**20% Glucose** (500 mL) – sterile filter

100g	Glucose

500 mL	sterile H_2_O

**1X enrichment media** (100 mL)

8% enrichment media	0.8% enrichment media

2X SC media (above)	50mL	50mL

20% glucose	40mL	4mL

sterile H_2_O	10mL	46mL

100 mg/mL Ampicillin	100μL	100μL

30 mg/mL Chloramphenicol	100μL	100μL

**YPD plates** (1 L)

10g	Yeast Extract

20g	Peptone

20g	Glucose

18g	Agar

1L	milliQ H_2_O
